# Measurement error associated with gait cycle selection in treadmill running at various speeds

**DOI:** 10.1101/2022.08.11.503696

**Authors:** Aaron S. Fox, Jason Bonacci, John Warmenhoven, Meghan F. Keast

**Affiliations:** Centre for Sport Research, School of Exercise and Nutrition Sciences, Deakin University, Geelong, Australia; School of Engineering and Information Technology, University of New South Wales, Canberra, Australia

## Abstract

A common approach in biomechanical analysis of running technique is to average data from several gait cycles to compute a ‘representative mean.’ However, the impact of the quantity and selection of gait cycles on biomechanical measures is not well understood. We examined the effects of gait cycle selection on kinematic data by: (i) comparing representative means calculated from varying numbers of gait cycles to ‘global’ means from the entire capture period; and (ii) comparing representative means from varying numbers of gait cycles sampled from different parts of the capture period. We used a public dataset (*n* = 28) of lower limb kinematics captured during a 30-second period of treadmill running at three speeds (2.5m · s^-1^, 3.5m · s^-1^ and 4.5m · s^-1^). ‘Ground truth’ values were determined by averaging data across all collected strides and compared to representative means calculated from random samples (1,000 samples) of *n* (range = 5—30) consecutive gait cycles. We also compared representative means calculated from n (range = 5—15) consecutive gait cycles randomly sampled (1,000 samples) from within the same data capture period. The mean, variance and range of the absolute error of the representative mean compared to the ‘ground truth’ mean progressively reduced across all speeds as the number of gait cycles used increased. Similar magnitudes of ‘error’ were observed between the 2.5m · s^-1^ and 3.5m · s^-1^ speeds at comparable gait cycle numbers — where the maximum errors were < 1.5 degrees even with a small number of gait cycles (i.e. 5-10). At the 4.5m · s^-1^ speed, maximum errors typically exceeded 2-4 degrees when a lower number of gait cycles were used. Subsequently, a higher number of gait cycles (i.e. 25-30) was required to achieve low errors (i.e. 1-2 degrees) at the 4.5m · s^-1^ speed. The mean, variance and range of absolute error of representative means calculated from different parts of the capture period was consistent irrespective of the number of gait cycles used. The error between representative means was low (i.e. <1.5 degrees) and consistent across the different number of gait cycles at the 2.5m · s^-1^ and 3.5m · s^-1^ speeds, and consistent but larger (i.e. up to 2-4 degrees) at the 4.5m · s^-1^ speed. Our findings suggest that selecting as many gait cycles as possible from a treadmill running bout will minimise potential ‘error.’ Analysing a small sample (i.e. 5-10 cycles) will typically result in minimal ‘error’ (i.e. < 2 degrees), particularly at lower speeds (i.e. 2.5m · s^-1^ and 3.5m · s^-1^). Researchers and clinicians should consider the balance between practicalities of collecting and analysing a smaller number of gait cycles against the potential ‘error’ when determining their methodological approach. Irrespective of the number of gait cycles used, we recommend that the potential ‘error’ introduced by the choice of gait cycle number be considered when interpreting the magnitude of effects in treadmill-based running studies.

## Introduction

Collecting and analysing biomechanical data is frequently used to examine running technique. A common methodological approach is to average data from several gait cycles to compute a given biomechanical measure. Calculating this ‘representative mean’ is thought to be representative of the individuals broader running technique. Given the inherent variability in human movement [1], the quantity and selection of gait cycles used to create this ‘representative mean’ appears an important choice in accurately quantifying an individuals running gait. However, the number of gait cycles used in biomechanical studies of running varies across the literature [2]. Further, very rarely (if ever) is the decision process underpinning the quantity and selection of gait cycles explained.

It is possible to collect a large number of gait cycles during biomechanical testing, especially during treadmill running. Enabling a participant to settle into a steady gait rhythm may better represent a habitual running pattern. While the collection of a large number of gait cycles can be relatively easy, it is important to give consideration to the analysis of this data. Inflated data cleaning (e.g. labelling and gap filling motion capture data) and analysis (e.g. processing frames via inverse kinematics) time occur when processing a running trial that uses many gait cycles. Similarly, trials with many gait cycles require greater data storage access due to larger file sizes. An additional consideration is which gait cycles are selected from within a capture period. Studies often perform an extended capture period where additional gait cycles are collected around those used for analysis (e.g. [4]). The impact of this gait cycle selection on biomechanical outcome measures is yet to be investigated. Better understanding of the impact of gait cycle selection on biomechanical outcome measures may help optimise data collection and analysis practices.

Oliveira and Pirscoveanu [2] examined the typical number of gait cycles used in running biomechanics studies. On average, 12 gait cycles were used to generate biomechanical outcome measures, though very few of these studies (5 out of 56) used more than 10 cycles [2]. The impact of sample size (i.e. 10 to 40 runners) and number of gait cycles (i.e. 5 to 40 steps) used on biomechanical outcome measures (i.e. foot contact time, loading rate, peak vertical ground reaction force, peak braking force, running speed, and foot contact angle) was also examined [2]. The authors found that greater than 10 strides are typically required to achieve stable biomechanical measures in runners and collecting at least 25 strides will increase the likelihood of achieving stability in the range of biomechanical measures examined [2]. These findings are specific to overground running and the set of biomechanical measures analysed. Treadmill running is often used in research [5], and it is plausible that the required number of gait cycles required to achieve stability may be different to overground running. Further, Oliveira and Pirscoveanu [2] did not examine lower limb kinematic variables commonly reported in gait biomechanics studies. These kinematic variables can be presented as both ‘zero-dimensional’ (0D; e.g. peak values) and ‘one-dimensional’ (1D; e.g. time-normalised kinematic waveform) variables [6]. Analyses of these common kinematic variables in both their 0D and 1D forms may provide valuable insight into the number of gait cycles required in biomechanical research. Lastly, Oliveira and Pirscoveanu’s [2] analyses were driven by understanding data stability and statistical significance between two running conditions (i.e. ‘normal’ vs. ‘silent’ running). A different approach focused on understanding the magnitude of ‘error’ introduced by analysing different numbers of gait cycles can further our understanding of how gait cycle selection practices impact biomechanical outcome measures. Specifically, understanding the potential ‘error’ introduced by selecting a different number of gait cycles can aid in interpreting the legitimacy of an effect (i.e. could small effects be due to the set of gait cycles selected).

We sought to extend our current understanding of how the quantity and selection of gait cycles impact lower limb kinematic measures from a 30-second data capture period of treadmill running. First, we examined the magnitude of ‘error’ introduced in the representative mean compared to the entire bout of treadmill running when the number of gait cycle samples is varied. Second, we examined the potential variation introduced in the representative mean when sampling a set number of gait cycles from different parts of the capture period.

## Methods

### Dataset

We used the public dataset of treadmill running biomechanics from Fukuchi et al. [7]. The specifics of this dataset can be found in the associated paper [7]. Briefly, this dataset contains lower-extremity kinematics and kinetics of 28 regular runners (27 male, 1 female; age = 34.8 ± 6.7 years; height = 176.0 ± 6.8 cm; mass = 69.6 ± 7.7 kg; running experience = 8.5 ± 7.0 years; running pace = 4.1 ± 0.4 min/km) [7]. Running kinematics were collected using a 12-camera 3D motion capture system (Raptor-4, Motion Analysis, Santa Rosa, CA, United States) and ground reaction force (GRF) data via an instrumented dual-belt treadmill (FIT, Bertec, Columbus, OH, United States) [7]. Participants ran on the treadmill at three speeds in order (2.5m · s^-1^, 3.5m · s^-1^ and 4.5m · s^-1^), during which a three-minute accommodation period was provided followed by a 30-second data collection period [7].

We processed the raw experimental data from Fukuchi et al. [7] using OpenSim 4.0 [8]. Segment geometry of a generic musculoskeletal model of the pelvis and lower limb provided by Lai et al. [9] were scaled for each participant using their static calibration trial, which was also used as a reference for adjusting marker positions on the model. Lower limb joint angles were calculated using filtered (10Hz low-pass 4^th^ order Butterworth) marker trajectory data within inverse kinematics analysis. GRF data were filtered using the same cut-off frequency and filter. The filtering procedures reflected those originally performed by Fukuchi et al. [7]. Foot strike and toe-off events were determined when the vertical GRF crossed a 20N threshold, also in line with the original work [7].

### Data Analysis

Kinematic variables common to gait biomechanics studies (i.e. hip flexion/extension, hip adduction/abduction, hip internal/external rotation, knee flexion and ankle plantarflexion/dorsiflexion) were extracted from the right limb for all participants. Data between consecutive foot strikes were extracted and time-normalised to 0-100% of the gait cycle. The time-normalised 1D curves were used in subsequent 1D analyses, while a set of peak variables (hip flexion, hip adduction, hip internal rotation, knee flexion, ankle dorsiflexion) were calculated and extracted for the 0D analyses.

To examine how the number of gait cycles used impacts the representative kinematic mean (i.e. aim 1), we determined ‘ground truth’ values to compare to for the 0D and 1D kinematic variables by calculating the mean from all available gait cycles in the 30-second capture period of treadmill running. This value was thought to be the ‘most representative’ of each participants average running kinematics and was not influenced by the selection of a subset of gait cycles. We then iteratively calculated mean values across the kinematic variables using a range (*n* = 5 — 30) of gait cycles from the data capture period. For each iteration, a random sample of *n* consecutive gait cycles were extracted and used to calculate a representative kinematic mean. We then compared this representative kinematic mean to the ‘ground truth’ value for the respective variable to determine the ‘error’ that gait cycle number selection could introduce.

To examine how sampling gait cycles from different sections of the capture period impacts the representative kinematic mean (i.e. aim 2), we iteratively calculated representative kinematic means using a range (*n* = 5 — 15) of randomly sampled consecutive gait cycles from different parts of the capture period. A smaller range of gait cycles was required for this analysis to avoid sharing gait cycles between the calculated means. For each sampling iteration, we randomly sampled *n* consecutive gait cycles from two non-overlapping parts of the capture period. We then compared the calculated representative kinematic means between the two parts to determine the ‘error’ or variation that selection of gait cycles from different parts of the capture period could introduce.

We quantified ‘error’ in a similar fashion across both aims. For 0D variables, the absolute difference between the representative mean and ‘ground truth’ (i.e. aim 1) or two representative means (i.e. aim 2) was recorded in each sampling iteration. For 1D variables, the absolute difference between the representative mean and ‘ground truth’ (i.e. aim 1) or two representative means (i.e. aim 2) at each point across the time-normalised gait cycle were calculated, and the peak difference recorded. The random sampling process for each *n* of gait cycles was repeated 1,000 times for each participant at each running speed — and the ‘error’ values collated to present descriptive statistics (i.e. mean ± standard deviation [SD], median, range, inter-quartile range) for each gait cycle number across the kinematic variables and running speeds.

## Results

### How does the number of gait cycles used impact the representative kinematic mean?

For the peak 0D kinematic variables, the mean, variance and range of the absolute error of the representative kinematic mean compared to the ‘ground truth’ mean progressively reduced as the number of gait cycles used increased (see Figures 1, 2 and 3). Similar magnitudes of ‘error’ were observed between the 2.5m · s^-1^ and 3.5m · s^-1^ speeds across the 0D kinematic variables at comparable gait cycle numbers — where the maximum errors were less than 1 degree even when using a small number of gait cycles. The maximum errors at the 4.5m · s^-1^ speed typically exceeded 1-2 degrees, particularly for peak hip and knee joint angles when a lower number of gait cycles were used. Subsequently, a much higher number of gait cycles (i.e. 25-30) were required at 4.5m · s^-1^ to achieve a similar magnitude of error seen at the slower running speeds. The larger ‘error’ values observed at 4.5m · s^-1^ were driven by a bimodal distribution — whereby certain sampling iterations within the same biomechanical measure could produce relatively higher versus lower errors (see Figure 3). The exception to this difference at the higher speed was for peak ankle dorsiflexion, where similarly low ‘error’ values and ranges (i.e. < 0.5 degrees) were observed across all speeds.

**Figure 1:**
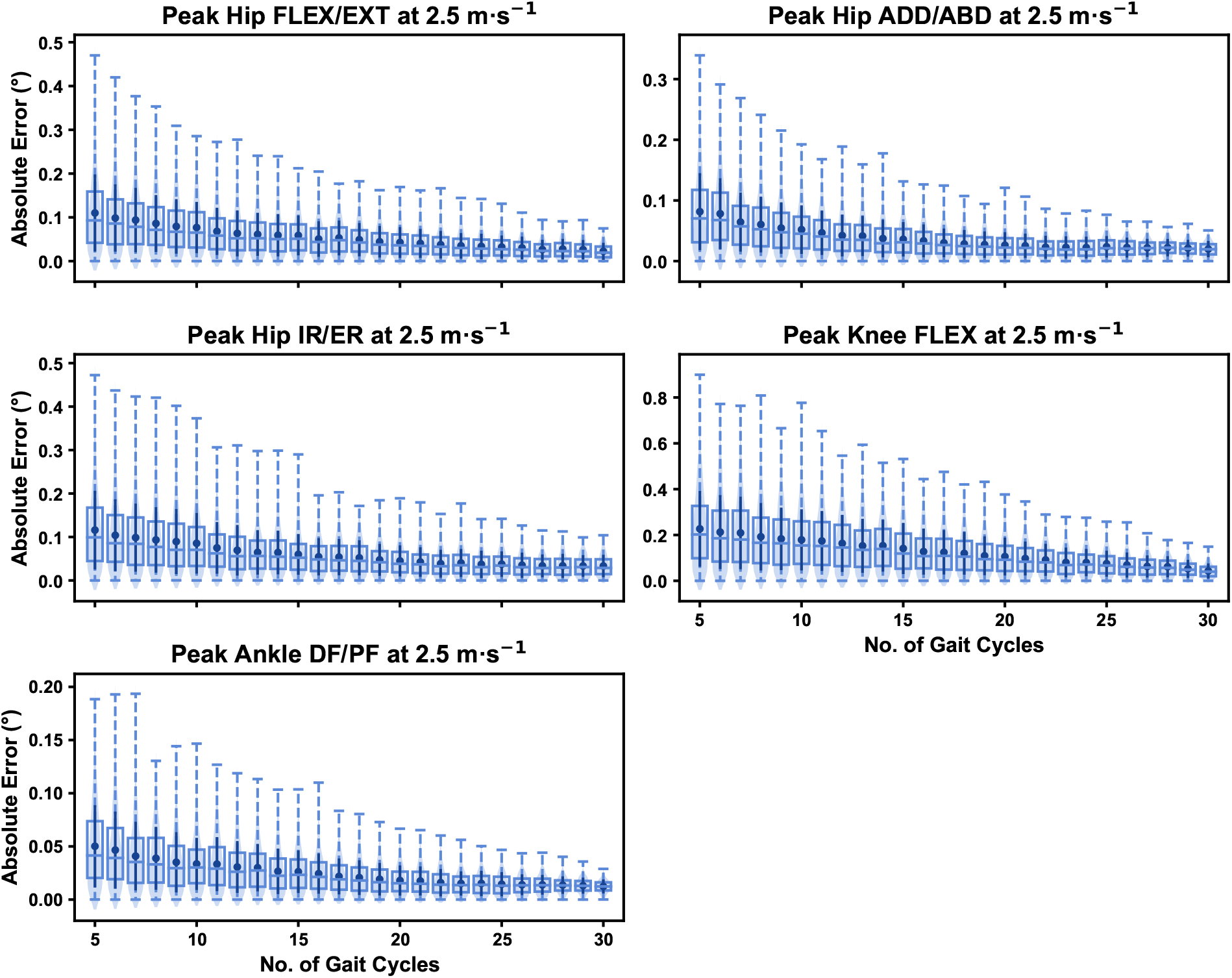
Absolute error in peak kinematic variables (i.e. zero-dimensional [0D]) when running at 2.5m · s^-1^ using a subset of gait cycles versus all gait cycles from the 30-second treadmill bout. Darker points and solid lines equate to the mean ± standard deviation. Horizontal lines within boxes equate to the median value, boxes indicate the 25*^th^* to 75*^th^* percentile, and dashed whiskers indicate the range. Shaded violins are included to illustrate the distribution of values. FLEX — flexion; EXT — extension; ADD — adduction; ABD — abduction; IR — internal rotation; ER — external rotation; DF — dorsiflexion; PF — plantarflexion.

**Figure 2:**
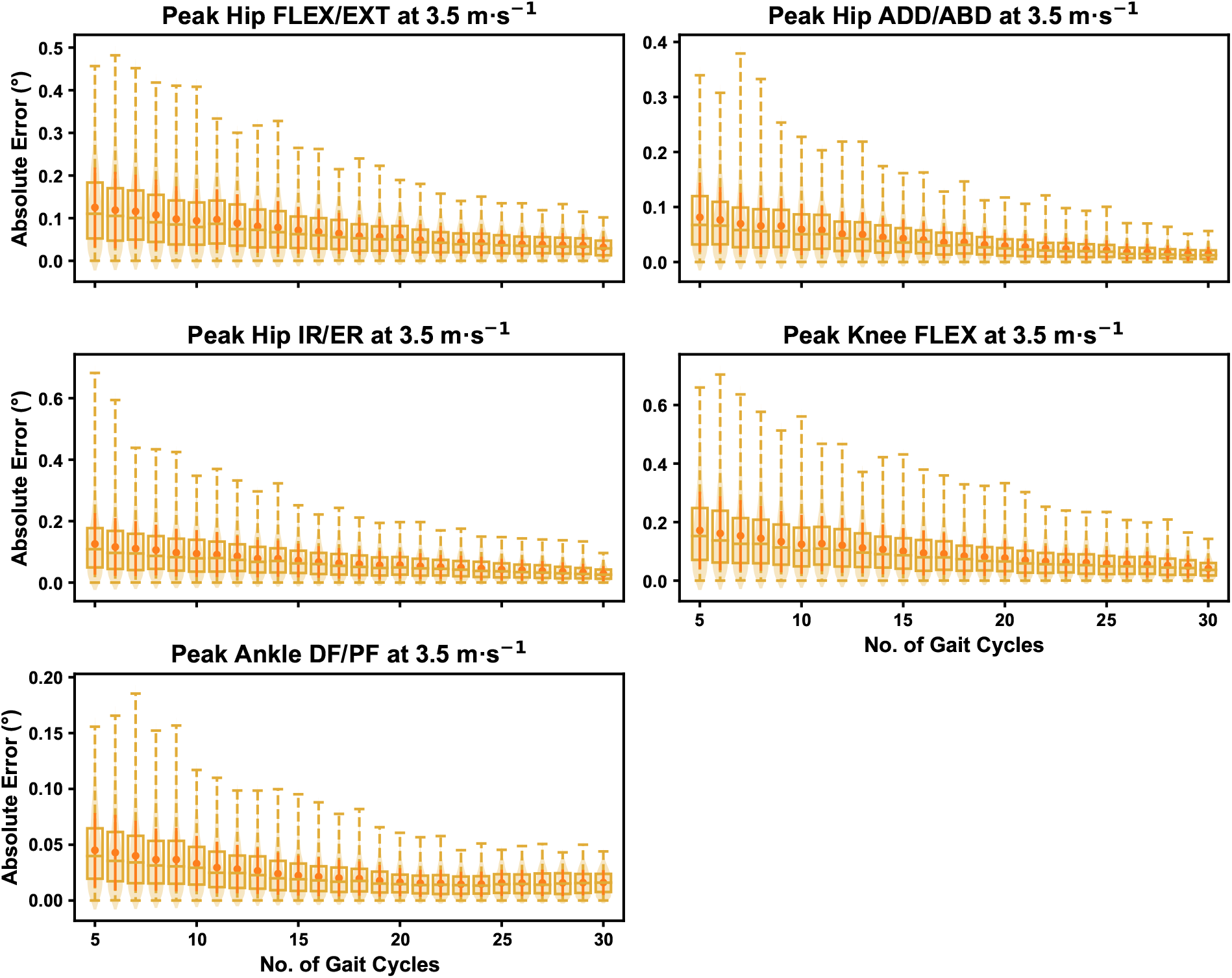
Absolute error in peak kinematic variables (i.e. zero-dimensional [0D]) when running at 3.5m · s^-1^ using a subset of gait cycles versus all gait cycles from the 30-second treadmill bout. Darker points and solid lines equate to the mean ± standard deviation. Horizontal lines within boxes equate to the median value, boxes indicate the 25*^th^* to 75*^th^* percentile, and dashed whiskers indicate the range. Shaded violins are included to illustrate the distribution of values. FLEX — flexion; EXT — extension; ADD — adduction; ABD — abduction; IR — internal rotation; ER — external rotation; DF — dorsiflexion; PF — plantarflexion.

**Figure 3:**
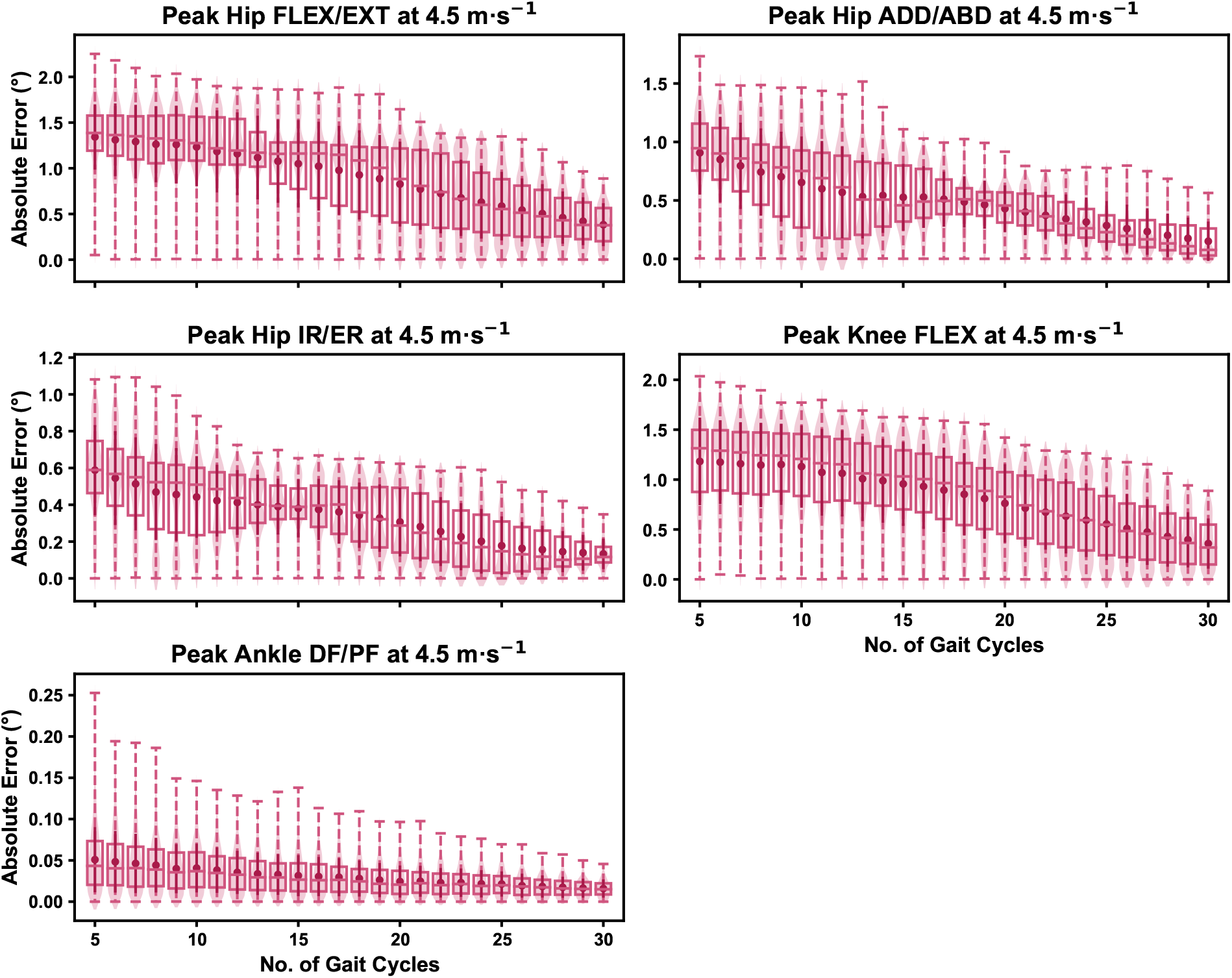
Absolute error in peak kinematic variables (i.e. zero-dimensional [0D]) when running at 4.5m · s^-1^ using a subset of gait cycles versus all gait cycles from the 30-second treadmill bout. Darker points and solid lines equate to the mean ± standard deviation. Horizontal lines within boxes equate to the median value, boxes indicate the 25*^th^* to 75*^th^* percentile, and dashed whiskers indicate the range. Shaded violins are included to illustrate the distribution of values. FLEX — flexion; EXT — extension; ADD — adduction; ABD — abduction; IR — internal rotation; ER — external rotation; DF — dorsiflexion; PF — plantarflexion.

We observed near identical characteristics of the mean, variance and range of the peak absolute error of the representative kinematic mean compared to the mean from all gait cycles for the 1D kinematic variables (see Figures 4, 5 and 6). As with the 0D variables, the potential ‘error’ reduced as the number of gait cycles increased, and similarly low magnitudes of ‘error’ (i.e. < 1 degree) were at the 2.5m · s^-1^ and 3.5m · s^-1^ speeds. Larger ‘errors’ exceeding 1-2 degrees with lower gait cycle numbers were present at the 4.5m · s^-1^ speed (with the exception of ankle dorsi/plantarflexion), with this again driven by a more bimodal distribution of samples (see Figure 6).

**Figure 4:**
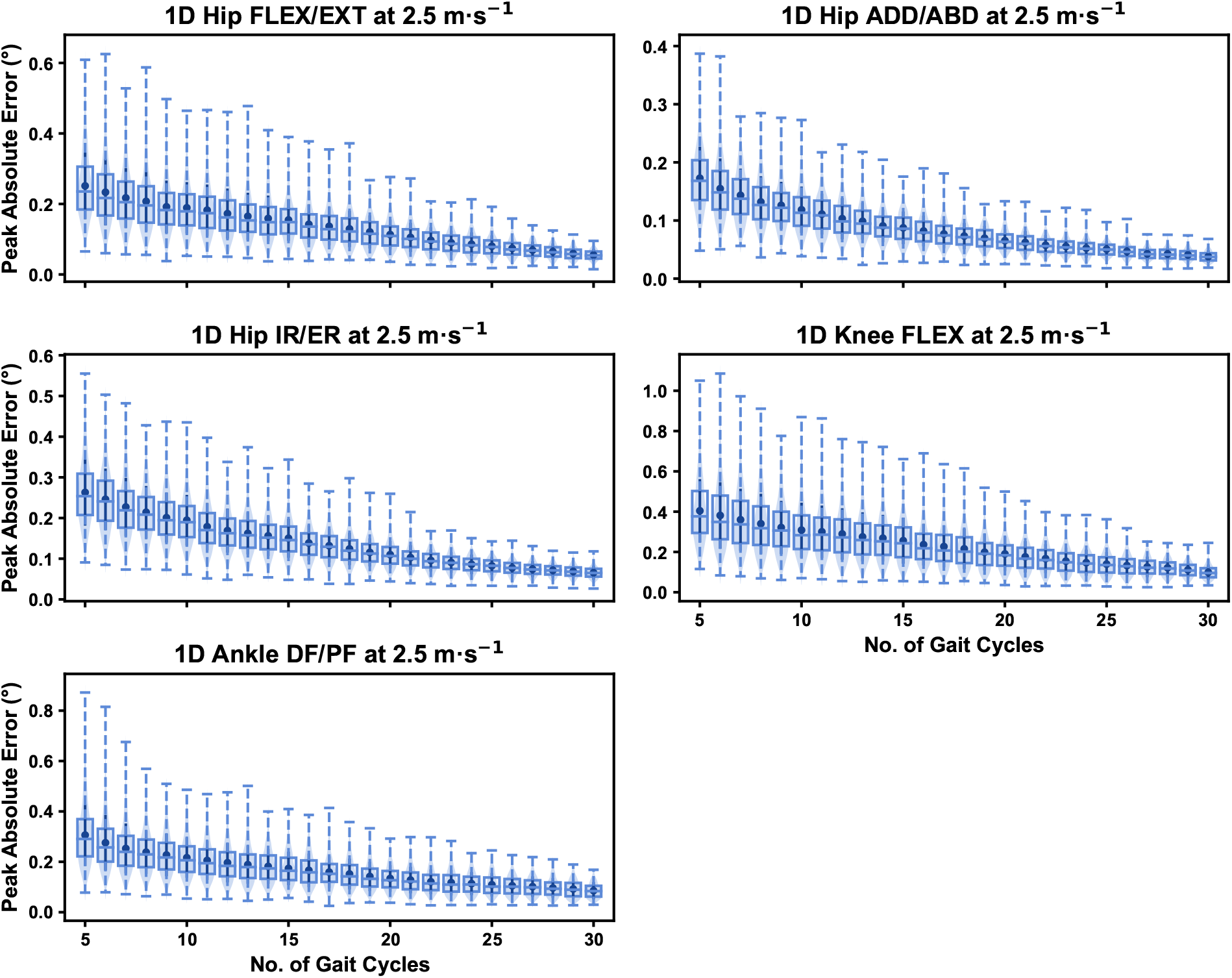
Peak absolute error in kinematic variables across the gait cycle (i.e. one-dimensional [1D]) when running at 2.5m · s^-1^ using a subset of gait cycles versus all gait cycles from the 30-second treadmill bout. Darker points and solid lines equate to the mean ± standard deviation. Horizontal lines within boxes equate to the median value, boxes indicate the 25*^th^* to 75*^th^* percentile, and dashed whiskers indicate the range. Shaded violins are included to illustrate the distribution of values. FLEX — flexion; EXT — extension; ADD — adduction; ABD — abduction; IR — internal rotation; ER — external rotation; DF — dorsiflexion; PF — plantarflexion.

**Figure 5:**
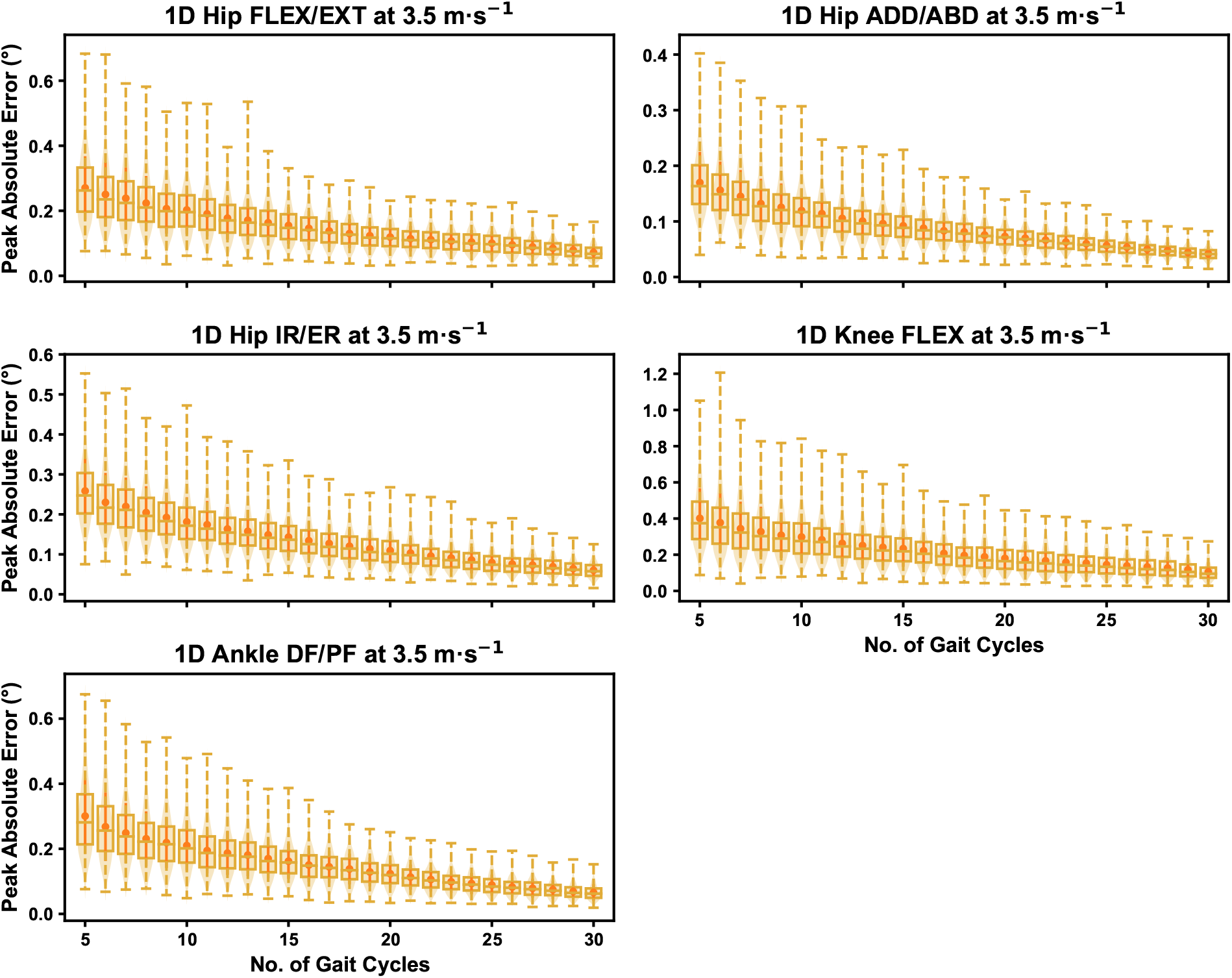
Peak absolute error in kinematic variables across the gait cycle (i.e. one-dimensional [1D]) when running at 3.5m · s^-1^ using a subset of gait cycles versus all gait cycles from the 30-second treadmill bout. Darker points and solid lines equate to the mean ± standard deviation. Horizontal lines within boxes equate to the median value, boxes indicate the 25*^th^* to 75*^th^* percentile, and dashed whiskers indicate the range. Shaded violins are included to illustrate the distribution of values. FLEX — flexion; EXT — extension; ADD — adduction; ABD — abduction; IR — internal rotation; ER — external rotation; DF — dorsiflexion; PF — plantarflexion.

**Figure 6:**
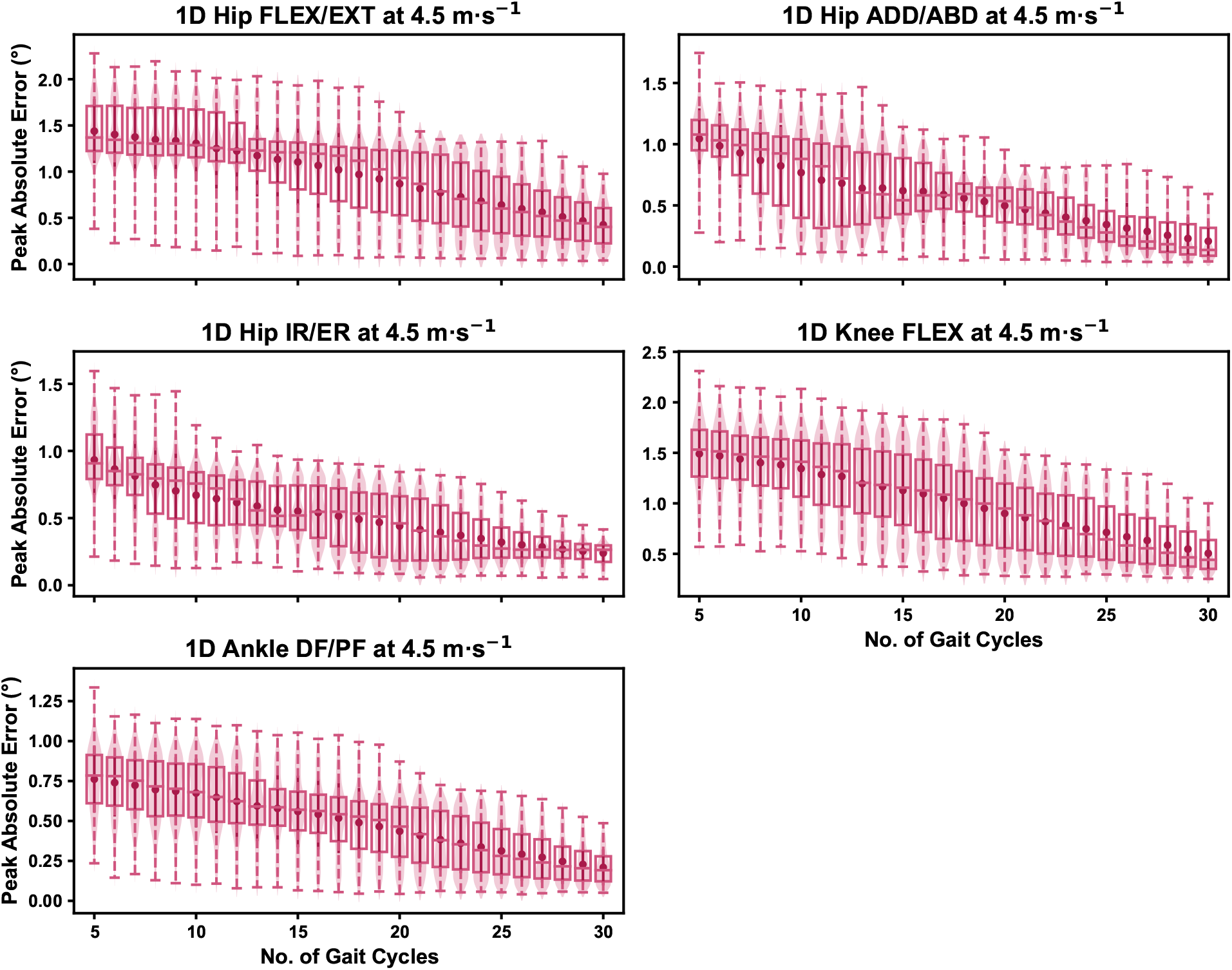
Peak absolute error in kinematic variables across the gait cycle (i.e. one-dimensional [1D]) when running at 4.5m · s^-1^ using a subset of gait cycles versus all gait cycles from the 30-second treadmill bout. Darker points and solid lines equate to the mean ± standard deviation. Horizontal lines within boxes equate to the median value, boxes indicate the 25*^th^* to 75*^th^* percentile, and dashed whiskers indicate the range. Shaded violins are included to illustrate the distribution of values. FLEX — flexion; EXT — extension; ADD — adduction; ABD — abduction; IR — internal rotation; ER — external rotation; DF — dorsiflexion; PF — plantarflexion.

### How does the selection of gait cycles impact the representative kinematic mean?

The mean, variance and range of the absolute error (or variation) of the representative kinematic mean compared to the mean from all gait cycles for the peak 0D kinematic variables remained relatively consistent irrespective of the number of gait cycles used when sampling from different parts of the capture period (see Figures 7, 8 and 9). At the 2.5m · s^-1^ and 3.5m · s^-1^ speeds, the variation in peak kinematic variables was always less than 1.5 degrees. However, peak knee flexion had the potential for larger variation compared to the remaining kinematic variables (see Figures 7 and 8). While the potential variation between gait cycle samples was consistent with increasing gait cycle numbers at the 4.5m · s^-1^ speed, a higher mean and range of potential variation (i.e. up to 2-4 degrees) was evident across the peak kinematic variables (with the exception of peak ankle dorsiflexion). As in the previous analyses, we observed a bimodal distribution of the samples at the 4.5m · s^-1^ speed (see Figure 9).

**Figure 7:**
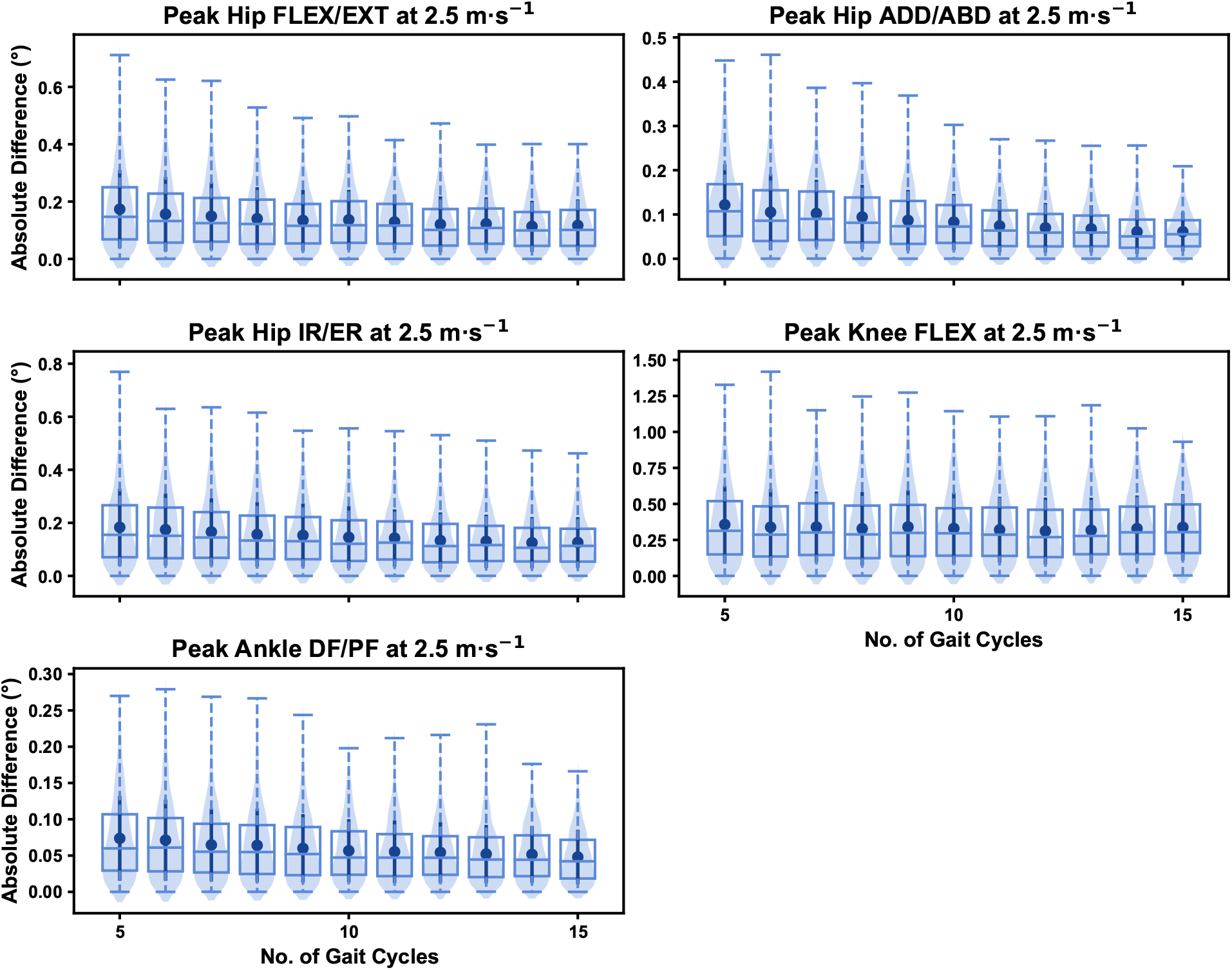
Absolute error in peak kinematic variables (i.e. zero-dimensional [0D]) when running at 2.5m · s^-1^ using a two comparative subsets of gait cycles from the 30-second treadmill bout. Darker points and solid lines equate to the mean ± standard deviation. Horizontal lines within boxes equate to the median value, boxes indicate the 25*^th^* to 75*^th^* percentile, and dashed whiskers indicate the range. Shaded violins are included to illustrate the distribution of values. FLEX — flexion; EXT — extension; ADD — adduction; ABD — abduction; IR — internal rotation; ER — external rotation; DF — dorsiflexion; PF — plantarflexion.

**Figure 8:**
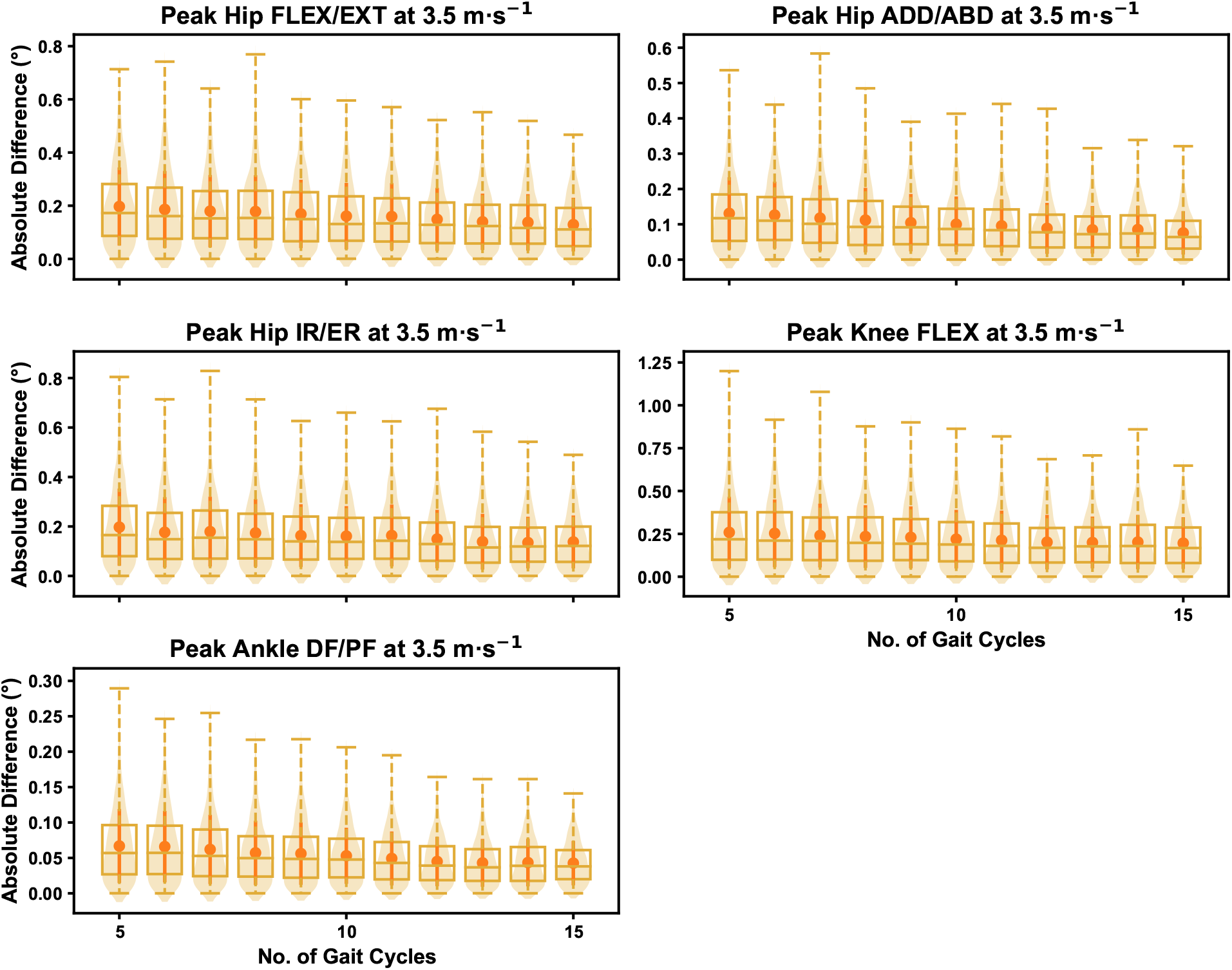
Absolute error in peak kinematic variables (i.e. zero-dimensional [0D]) when running at 3.5m · s^-1^ using a two comparative subsets of gait cycles from the 30-second treadmill bout. Darker points and solid lines equate to the mean ± standard deviation. Horizontal lines within boxes equate to the median value, boxes indicate the 25*^th^* to 75*^th^* percentile, and dashed whiskers indicate the range. Shaded violins are included to illustrate the distribution of values. FLEX — flexion; EXT — extension; ADD — adduction; ABD — abduction; IR — internal rotation; ER — external rotation; DF — dorsiflexion; PF — plantarflexion.

**Figure 9:**
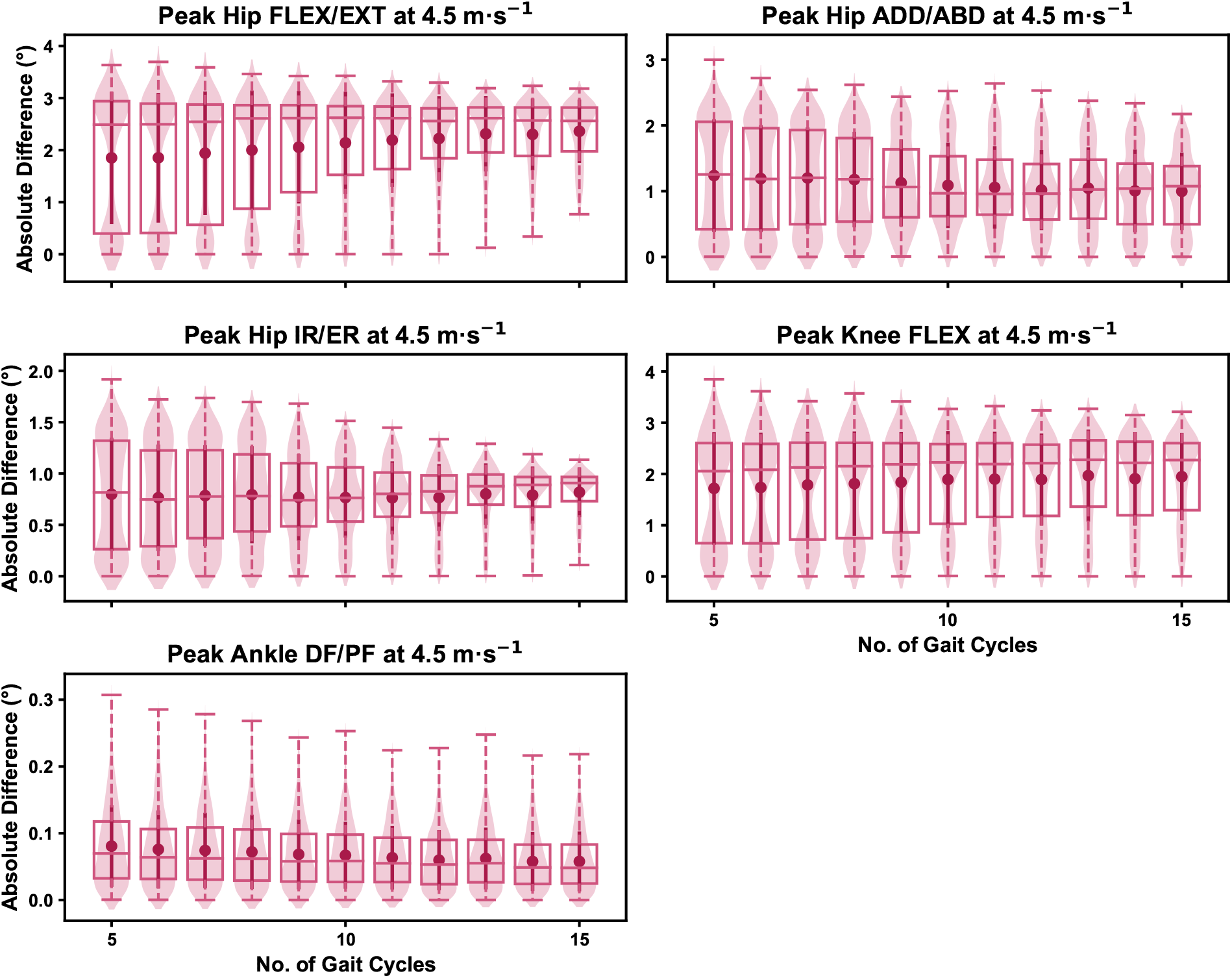
Absolute error in peak kinematic variables (i.e. zero-dimensional [0D]) when running at 4.5m · s^-1^ using a two comparative subsets of gait cycles from the 30-second treadmill bout. Darker points and solid lines equate to the mean ± standard deviation. Horizontal lines within boxes equate to the median value, boxes indicate the 25*^th^* to 75*^th^* percentile, and dashed whiskers indicate the range. Shaded violins are included to illustrate the distribution of values. FLEX — flexion; EXT — extension; ADD — adduction; ABD — abduction; IR — internal rotation; ER — external rotation; DF — dorsiflexion; PF — plantarflexion.

We observed similar characteristics for the mean, variance and range of the absolute error (or variation) of the representative kinematic mean compared to the mean from all gait cycles for the 1D kinematic variables when sampling gait cycles from different parts of the capture period (see Figures 10, 11 and 12). The potential variation remained low (i.e. < 1.5 degrees) and consistent across the different number of gait cycles at the 2.5m · s^-1^ and 3.5m · s^-1^ speeds (see Figures 10 and 11). The potential variation remained consistent but increased in magnitude (i.e. up to 2-4 degrees), and shifted to a bimodal distribution at the 4.5m · s^-1^ speed (see Figure 12). In contrast to the 0D variables, this shift was evident in all 1D kinematic variables (including ankle dorsi/plantarflexion).

**Figure 10:**
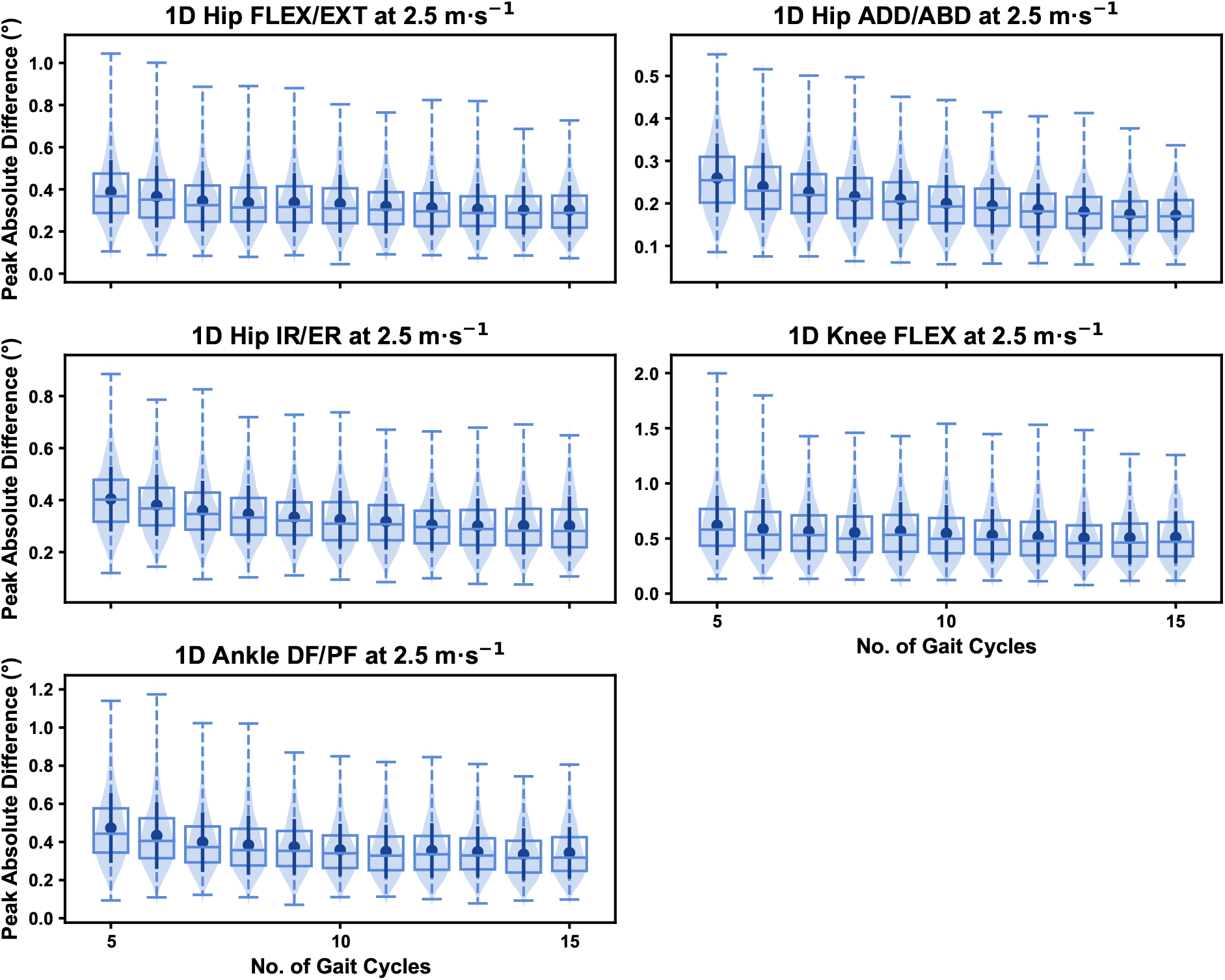
Peak absolute error in kinematic variables across the gait cycle (i.e. one-dimensional [1D]) when running at 2.5m · s^-1^ using two comparative subsets of gait cycles from the 30-second treadmill bout. Darker points and solid lines equate to the mean ± standard deviation. Horizontal lines within boxes equate to the median value, boxes indicate the 25*^th^* to 75*^th^* percentile, and dashed whiskers indicate the range. Shaded violins are included to illustrate the distribution of values. FLEX — flexion; EXT — extension; ADD — adduction; ABD — abduction; IR — internal rotation; ER — external rotation; DF — dorsiflexion; PF — plantarflexion.

**Figure 11:**
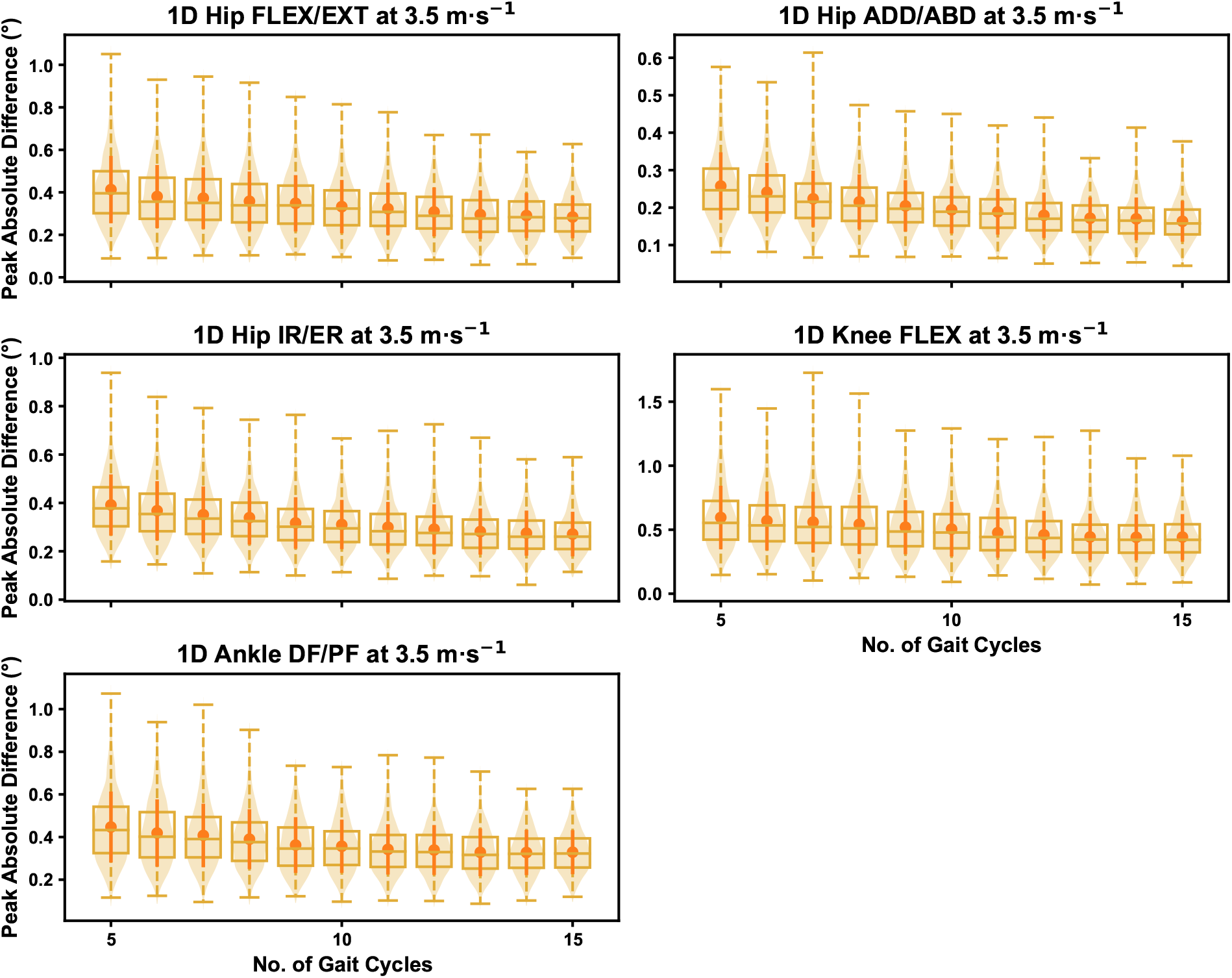
Peak absolute error in kinematic variables across the gait cycle (i.e. one-dimensional [1D]) when running at 3.5m · s^-1^ using two comparative subsets of gait cycles from the 30-second treadmill bout. Darker points and solid lines equate to the mean ± standard deviation. Horizontal lines within boxes equate to the median value, boxes indicate the 25*^th^* to 75*^th^* percentile, and dashed whiskers indicate the range. Shaded violins are included to illustrate the distribution of values. FLEX — flexion; EXT — extension; ADD — adduction; ABD — abduction; IR — internal rotation; ER — external rotation; DF — dorsiflexion; PF — plantarflexion.

**Figure 12:**
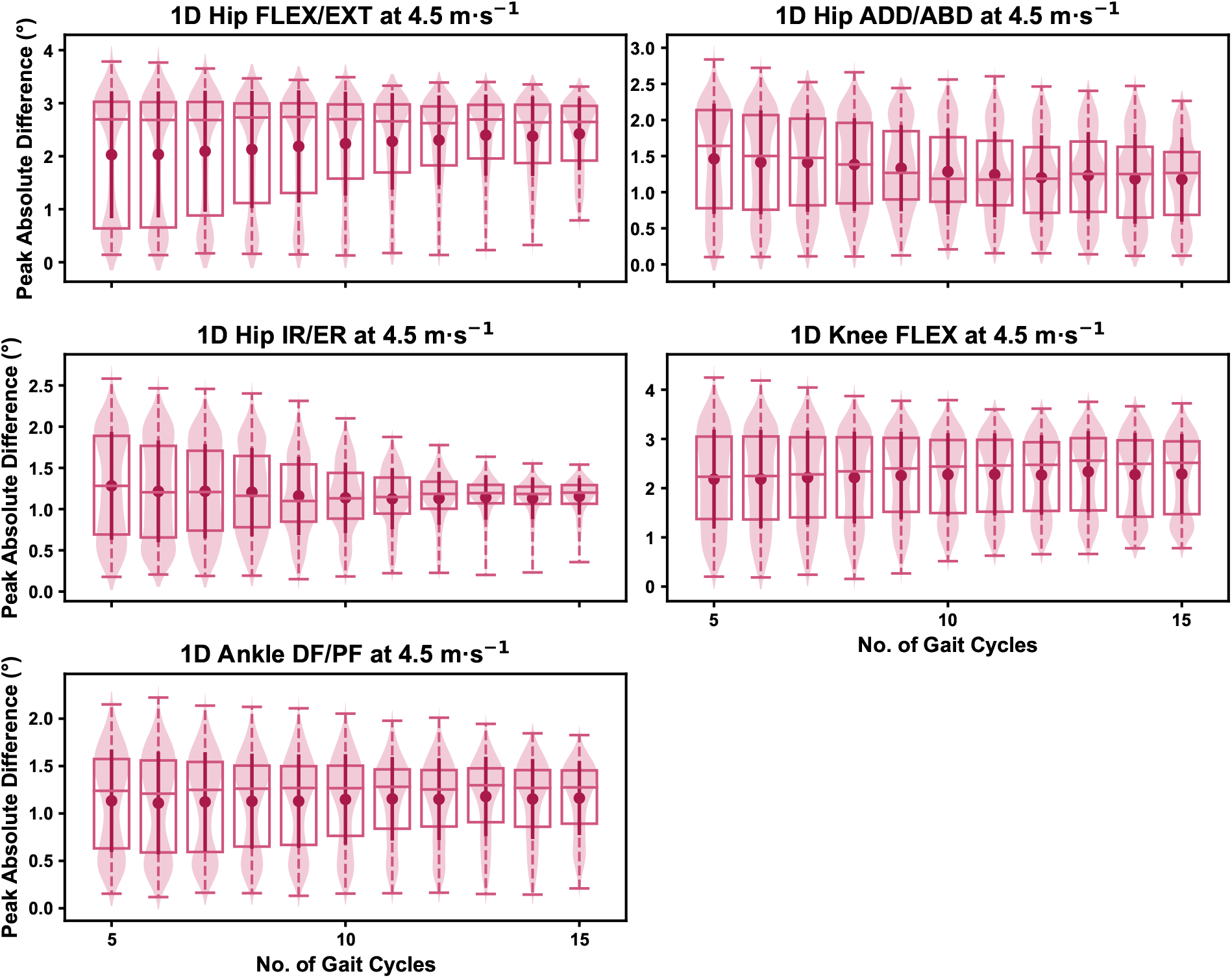
Peak absolute error in kinematic variables across the gait cycle (i.e. one-dimensional [1D]) when running at 4.5m · s^-1^ using two comparative subsets of gait cycles from the 30-second treadmill bout. Darker points and solid lines equate to the mean ± standard deviation. Horizontal lines within boxes equate to the median value, boxes indicate the 25*^th^* to 75*^th^* percentile, and dashed whiskers indicate the range. Shaded violins are included to illustrate the distribution of values. FLEX — flexion; EXT — extension; ADD — adduction; ABD — abduction; IR — internal rotation; ER — external rotation; DF — dorsiflexion; PF — plantarflexion.

## Discussion

Biomechanical studies of running often use a subset of gait cycles from a running bout or capture period, and average across these cycles to calculate an individual’s representative mean. We examined the impact of the quantity and selection of gait cycles from within the a capture period on the magnitude of ‘error’ in lower limb kinematic measures during a continuous bout of treadmill running. We found that including a greater number of gait cycles to calculate the representative kinematic mean reduces the magnitude and range of potential ‘error.’ The potential error using a small number of gait cycles (i.e. *n* = 5-10) was low (i.e. typically < 1 degree) when running at 2.5m · s^-1^ and 3.5m · s^-1^, and hence we noted an effect of diminishing returns (i.e. limited improvement in error reduction above 5-10 gait cycles) by including more gait cycles at these slower speeds. Using a similarly low number of gait cycles did slightly inflate potential ‘error’ (i.e. 1-4 degrees) when running at 4.5m · s^-1^. We also found small magnitudes of ‘error’ in representative kinematic means across all running speeds when selecting gait cycles from different parts of the capture period, and these remained relatively consistent irrespective of the number of gait cycles used.

We found that the ‘error’ between the representative kinematic means and the associated ‘ground truth’ values progressively reduced with an increasing number of gait cycles. Using a greater number of gait cycles equated to using a higher proportion of data that were used to create the ‘ground truth’ — hence this result is not surprising. More noteworthy is the scale of ‘error’ when using a reduced number of gait cycles (i.e. *n* = 5-10) and the diminishing effect of using a larger number (i.e. *n* > 15) of gait cycles. We typically observed that the maximum ‘error’ or variation with respect to the ‘ground truth’ was less than one degree, even at the lowest number of gait cycles used when running at the 2.5m · s^-1^ and 3.5m · s^-1^. This error increased up to three degrees at 4.5m · s^-1^. Reducing the potential ‘error’ compared to the ground truth appeared to be the main effect of increasing the number of gait cycles used. However, the reduction in potential error typically plateaued and a diminished benefit observed when using above 15-20 gait cycles. These patterns were consistent across both the 0D and 1D kinematic data. The notion of diminishing returns above 15-20 gait cycles contrasts with the findings of Oliveira and Pirscoveanu [2] — whereby data stability was not achieved in most runners using this number of gait cycles. Clear differences between our study and this existing work [2] were the metrics used to define ‘error’ or stability, the biomechanical measures analysed (i.e. joint kinematics vs. mostly kinetic variables), and the use of treadmill (including a 3-minute familiarisation period) versus overground running. The latter may represent an important distinction, whereby the familiarisation period combined with the more continuous approach of treadmill running led to participants settling into a more stable rhythm during the data capture period. Forrester [10] performed a series of simulations using a similar sequential analysis technique to Oliveira and Pirscoveanu [2] to determine the number of trials required for biomechanical measures with generic means and standard deviations. This work [10] proposed that nine (± 8) trials were required to achieve stability of the mean, which is more in line with our findings of diminishing returns in ‘error’ at 15-20 gait cycles. The mean and variation of the biomechanical outcome measure being examined likely plays a role in the potential ‘error.’ We saw the largest potential ‘errors’ in hip and knee flexion when using a smaller number of gait cycles and this is not surprising given these measures had the largest means and standard deviations within the dataset [7].

Despite the potential for diminishing returns, our data suggests that researchers can minimise the potential ‘error’ in representative kinematic means by using more gait cycles. A simplistic recommendation from our analyses would be to use as many gait cycles as possible. However, this ignores the practical considerations of storing, cleaning and processing larger biomechanical data files. Certain circumstances, such as a large participant sample or timely computational measures (e.g. muscle forces derived from optimisation approaches), may make using 20+ gait cycles impractical. Our recommendation is to balance the practical considerations against the potential ‘error’ or variation in the data that can be tolerated. Consideration should be given to the accuracy of the measure, or size of the effect researchers or clinicians are interested in measuring. For example, using less than ten gait cycles to explore a small effect (i.e. < 1-2 degrees) in 1D hip or knee flexion continua may be unwise, as the potential variation in the calculated means could exceed the magnitude of the effect of interest. Our data suggests that the smaller the expected effect or magnitude of effect of interest, the greater number of gait cycles necessary for the analyses.

We observed relatively small variations (i.e. < 1.5 degrees) between representative kinematic means calculated from gait cycle samples extracted from different parts of the 2.5m · s^-1^ and 3.5m · s^-1^ capture periods, while these slightly increased (i.e. 2-4 degrees) when examining the 4.5m · s^-1^ speed. This magnitude of variation remained consistent irrespective of the total number of gait cycles used. These findings suggest that once the number of gait cycles used for analysis is selected, the selection of these from within a capture period will introduce a small, but consistent amount of ‘error.’ The inherent variability in human movement [1] is the likely and potentially unavoidable cause of this variation. We randomly sampled differing sections of the capture period as part of our analyses, and at-times this generated near zero variation between the two representative means. Without further inspection of our data, we cannot confirm what generated the reduced variation — but we hypothesise that the samples with minimal to no variation likely stemmed from using sections of the capture period in close proximity to one another. We also cannot determine which section of the running bout is more representative or ‘accurate,’ as we only compared between samples and did not extend this comparison to the ‘ground truth’ values. Our data can only be used to infer the potential magnitude of variation expected when using gait cycles from different parts of a 30-second capture period. The magnitude of this variation appears to be driven by the scale of the mean and standard deviation of the measure (i.e. kinematic measures with higher means and standard deviations incur a greater magnitude of variation). It is also plausible that greater variation could be seen during longer capture periods than that used in the present study (i.e. 30-seconds) or when comparing gait cycles from capture periods separated by a longer time period (e.g. two capture periods at either end of a 5+ minute running bout). The dataset we used did not allow for these analyses to be conducted, yet present relevant avenues for further research on this topic. The practical implications of these findings once again relate to the confidence we can have in measuring an effect on lower limb kinematics during treadmill running. If our observed effect does not exceed the typical variation seen when sampling from different parts of the capture period, there is a possibility that the observed effect is simply noise due to the gait cycles sampled.

Running at 4.5m · s^-1^ induced greater ‘error’ relative to the ‘ground truth’ and between representative means from different parts of the capture period compared to running at 2.5m · s^-1^ and 3.5m · s^-1^. There are various potential reasons for these results. Faster running speeds induce larger means and standard deviations across kinematic variables [7], particularly in hip and knee flexion where more dramatic increases in ‘error’ were observed. We propose that the larger means and standard deviations at higher speeds introduce a greater magnitude of variation across gait cycles. An increase in gait speed could also be considered a changed task constraint on the running movement [11], and this change in constraint could have affected the role and magnitude of variability at certain joints. Movement variability may help explain the greater potential for ‘error’ when sampling from different gait cycles at different running speeds. The changed task constraint, and greater kinematic means and standard deviations with increased running speed [7] may engender expectations of a consistent increase in potential ‘error’ with increased running speed. It is therefore surprising that the increase in ‘error’ or variation was inconsistent, and most evident and prominent when only when running at 4.5m · s^-1^ speed. Within the dataset examined, participants ran for a three minute accommodation period at each speed, following which data were collected over a 30 second period [7]. The order of running conditions (i.e. 2.5m · s^-1^, 3.5m · s^-1^, 4.5m · s^-1^) was kept consistent for each participant [7]. It is possible that these experimental procedures (i.e. running at the fastest speed towards the end of the running period) could have introduced some fatigue when running at 4.5m · s^-1^. Running in a fatigued state can increase biomechanical variability [13], while also altering running kinematics compared to a non-fatigued state [16]. If fatigue was present during the final bout of running, it could have resulted in greater kinematic variability or a change in running kinematics during the 30 second period of data collection. Alternatively, fatigue may have begun to set in within the final 30 seconds of the run — potentially inducing a change in running kinematics within the period where data were collected. This latter explanation may explain the bimodal distribution in ‘error’ we observed in the 4.5m · s^-1^ running bout, whereby larger ‘errors’ may have been observed with gait cycles from earlier versus later parts of the 30 second capture period. Given we did not explicitly consider the sections where gait cycles were sampled from, this notion is speculative. It should also be noted that studies examining changes in running biomechanics with fatigue [16] have used more intense and longer duration exercise protocols that what participants experienced in our study. Despite the lack of understanding around the potential mechanism, our study demonstrates a need to consider gait cycle sampling practices when running at faster speeds, and potentially when fatigue is present.

It is important to note that the measurement ‘error’ or variation based on gait cycle selection were small compared to other established sources of error during biomechanical data collection and analysis [28]. The magnitude of ‘error’ in the present study is eclipsed by the errors or variation introduced by soft-tissue artefact associated with skin-mounted markers [20], different joint coordinate systems [23] or gait models [24], kinematic algorithm choice [25], tester experience [27], or different measurement approaches (i.e. marker vs. marker-less) [28]. The number of gait cycles used for analysis is likely less important when considering the size of a measured effect against the potential ‘error’ or variation introduced by other methodological decisions. Future research should also better define practically meaningful effects for biomechanical outcome measures. Using similar methods to those in other fields for defining the smallest effect size of interest [29] may help inform whether the magnitude of errors are acceptable for practical use and interpretation.

Our results must be considered with respect to the limitations in our approach. We only examined conditions where *n* consecutive gait cycles were sampled from a 30-second capture period during a continuous bout of treadmill running at three set speeds. Different results might be expected with non-consecutive selection of samples from the capture period, or under different running conditions (e.g. outdoor overground running; slower or faster speeds). We also focused on peak and 1D waveform data of lower limb kinematic variables. Other biomechanical outcome measures (e.g. joint moments, estimates of muscle activation and forces) may incur variable magnitudes of ‘error’ or variation with respect to gait cycle selection. We inferred ‘error’ via comparison to values calculated from all gait cycles in the running bout (i.e. our ‘ground truth’ value). Although we deemed this the best approach within our study, it is important to acknowledge that these values may still not represent the individuals exact or true running kinematics. Lastly, we investigated kinematic measures at a univariate joint level. Our findings are therefore not applicable to studies examining covariance or dynamics across joints during gait.

## Conclusions

We identified the range of potential ‘error’ or variation in lower limb kinematics associated with the quantity and selection of gait cycles used from a data capture period of continuous treadmill running. Our findings suggest that including as many gait cycles as possible from the running bout will minimise ‘error.’ However, the error associated with only a small sample of gait cycles (i.e. 5-10 gait cycles) was typically quite small (< 3 degrees) when running at 2.5m · s^-1^ and 3.5m · s^-1^. Larger potential ‘errors’ or variation were observed when analysing kinematic variables with larger means and standard deviations, and when running at faster speeds (i.e. 4.5m · s^-1^). Researchers and clinicians should balance the benefits of a reduction in potential ‘error’ with the challenges of collecting, processing and analysing a large number of gait cycles when determining their methodological approach. We recommend that the potential ‘error’ or variation introduced by the quantity and selection of gait cycles be considered when interpreting effects from treadmill-based running studies. Specifically, researchers must consider the magnitude of potential ‘error’ against the identified effects between groups or following an intervention.

## Notes

### Competing Interest Statement

The authors have declared no competing interest.

## References

[1] Emmerik REA van, Wegen EEH van. On Variability and Stability in Human Movement. Journal of Applied Biomechanics 2000;16:394–406. doi:10.1123/jab.16.4.394.

[2] Oliveira AS, Pirscoveanu CI. Implications of sample size and acquired number of steps to investigate running biomechanics. Scientific Reports 2021;11:3083. doi:10.1038/s41598-021-82876-z.

[3] Fellin RE, Manal K, Davis IS. Comparison of lower extremity kinematic curves during overground and treadmill running. Journal of Applied Biomechanics 2010;26:407–14. doi:10.1126/scisignal.2001449.Engineering.

[4] Fox AS, Ferber R, Bonacci J. Kinematic and Coordination Variability in Individuals With Acute and Chronic Patellofemoral Pain. Journal of Applied Biomechanics 2021;37:463–70. doi:10.1123/jab.2020-0401.

[5] Van Hooren B, Fuller JT, Buckley JD, Miller JR, Sewell K, Rao G, et al. Is Motorized Treadmill Running Biomechanically Comparable to Overground Running? A Systematic Review and Meta-Analysis of Cross-Over Studies. Sports Medicine 2020;50:785–813. doi:10.1007/s40279-019-01237-z.

[6] Pataky TC, Vanrenterghem J, Robinson MA. The probability of false positives in zero-dimensional analyses of one-dimensional kinematic, force and EMG trajectories. Journal of Biomechanics 2016;49:1468–76. doi:10.1016/j.jbiomech.2016.03.032.

[7] Fukuchi RK, Fukuchi CA, Duarte M. A public dataset of running biomechanics and the effects of running speed on lower extremity kinematics and kinetics. PeerJ 2017;5:e3298. doi:10.7717/peerj.3298.

[8] Delp SL, Anderson FC, Arnold AS, Loan P, Habib A, John CT, et al. OpenSim: Open-Source Software to Create and Analyze Dynamic Simulations of Movement. IEEE Transactions on Biomedical Engineering 2007;54:1940–50. doi:10.1109/TBME.2007.901024.

[9] Lai AKM, Arnold AS, Wakeling JM. Why are Antagonist Muscles Co-activated in My Simulation? A Musculoskeletal Model for Analysing Human Locomotor Tasks. Annals of Biomedical Engineering 2017;45:2762–74. doi:10.1007/s10439-017-1920-7.

[10] Forrester SE. Selecting the number of trials in experimental biomechanics studies. International Biomechanics 2015;2:62–72. doi:10.1080/23335432.2015.1049296.

[11] Newell KM. Coordination, Control and Skill. Advances in Psychology 1985;27:295–317. doi:10.1016/S0166-4115(08)62541-8.

[12] Meardon SA, Hamill J, Derrick TR. Running injury and stride time variability over a prolonged run. Gait & Posture 2011;33:36–40. doi:10.1016/j.gaitpost.2010.09.020.

[13] Chen TL-W, Wong DW-C, Wang Y, Tan Q, Lam W-K, Zhang M. Changes in segment coordination variability and the impacts of the lower limb across running mileages in half marathons: Implications for running injuries. Journal of Sport and Health Science 2022;11:67–74. doi:10.1016/j.jshs.2020.09.006.

[14] Bazuelo-Ruiz B, Durá-Gil JV, Palomares N, Medina E, Llana-Belloch S. Effect of fatigue and gender on kinematics and ground reaction forces variables in recreational runners. PeerJ 2018;6:e4489. doi:10.7717/peerj.4489.

[15] Derrick TR, Dereu D, McLean SP. Impacts and kinematic adjustments during an exhaustive run. Medicine and Science in Sports and Exercise 2002;34:998–1002. doi:10.1097/00005768-200206000-00015.

[16] Mizrahi J, Verbitsky O, Isakov E, Daily D. Effect of fatigue on leg kinematics and impact acceleration in long distance running. Human Movement Science 2000;19:139–51. doi:10.1016/S0167-9457(00)00013-0.

[17] Leardini A, Chiari L, Croce UD, Cappozzo A. Human movement analysis using stereophotogrammetry. Gait & Posture 2005;21:212–25. doi:10.1016/j.gaitpost.2004.05.002.

[18] Benoit DL, Damsgaard M, Andersen MS. Surface marker cluster translation, rotation, scaling and deformation: Their contribution to soft tissue artefact and impact on knee joint kinematics. Journal of Biomechanics 2015;48:2124–9. doi:10.1016/j.jbiomech.2015.02.050.

[19] Fiorentino NM, Atkins PR, Kutschke MJ, Goebel JM, Foreman KB, Anderson AE. Soft tissue artifact causes significant errors in the calculation of joint angles and range of motion at the hip. Gait & Posture 2017;55:184–90. doi:10.1016/j.gaitpost.2017.03.033.

[20] D’Isidoro F, Brockmann C, Ferguson SJ. Effects of the soft tissue artefact on the hip joint kinematics during unrestricted activities of daily living. Journal of Biomechanics 2020;104:109717. doi:10.1016/j.jbiomech.2020.109717.

[21] Baudet A, Morisset C, D’Athis P, Maillefert J-F, Casillas J-M, Ornetti P, et al. Cross-Talk Correction Method for Knee Kinematics in Gait Analysis Using Principal Component Analysis (PCA): A New Proposal. PLoS ONE 2014;9:e102098. doi:10.1371/journal.pone.0102098.

[22] Colle F, Lopomo N, Visani A, Zaffagnini S, Marcacci M. Comparison of three formal methods used to estimate the functional axis of rotation: an extensive in-vivo analysis performed on the knee joint. Computer Methods in Biomechanics and Biomedical Engineering 2016;19:484–92. doi:10.1080/10255842.2015.1042464.

[23] Sauret C, Pillet H, Skalli W, Sangeux M. On the use of knee functional calibration to determine the medio-lateral axis of the femur in gait analysis: Comparison with EOS biplanar radiographs as reference. Gait and Posture 2016;50:180–4. doi:10.1016/j.gaitpost.2016.09.008.

[24] Mentiplay BF, Clark RA. Modified conventional gait model versus cluster tracking: Test-retest reliability, agreement and impact of inverse kinematics with joint constraints on kinematic and kinetic data. Gait & Posture 2018;64:75–83. doi:10.1016/j.gaitpost.2018.05.033.

[25] Kainz H, Modenese L, Lloyd DG, Maine S, Walsh HPJ, Carty CP. Joint kinematic calculation based on clinical direct kinematic versus inverse kinematic gait models. Journal of Biomechanics 2016;49:1658–69. doi:10.1016/j.jbiomech.2016.03.052.

[26] Leigh RJ, Pohl MB, Ferber R. Does tester experience influence the reliability with which 3D gait kinematics are collected in healthy adults? Physical Therapy in Sport 2014;15:112–6. doi:10.1016/j.ptsp.2013.04.003.

[27] Sinclair J, Hebron J, Taylor PJ. The influence of tester experience on the reliability of 3D kinematic information during running. Gait & Posture 2014;40:707–11. doi:10.1016/j.gaitpost.2014.06.004.

[28] Ceseracciu E, Sawacha Z, Cobelli C. Comparison of Markerless and Marker-Based Motion Capture Technologies through Simultaneous Data Collection during Gait: Proof of Concept. PLoS ONE 2014;9:e87640. doi:10.1371/journal.pone.0087640.

[29] Lakens D, Scheel AM, Isager PM. Equivalence Testing for Psychological Research: A Tutorial. Advances in Methods and Practices in Psychological Science 2018;1:259–69. doi:10.1177/2515245918770963.

